# Mecamylamine inhibits seizure-like activity in CA1-CA3 hippocampus through antagonism to nicotinic receptors

**DOI:** 10.1101/2020.09.21.306019

**Authors:** Olha Zapukhliak, Olga Netsyk, Artur Romanov, Oleksandr Maximyuk, Murat Oz, Gregory L. Holmes, Oleg Krishtal, Dmytro Isaev

**Author notes:** Corresponding author (DS).

## Abstract

Cholinergic modulation of hippocampal network function is implicated in multiple behavioral and cognitive states. Activation of nicotinic and muscarinic acetylcholine receptors affects neuronal excitability, synaptic transmission and rhythmic oscillations in the hippocampus. In this work, we study the ability of the cholinergic system to sustain hippocampal epileptiform activity independently from glutamate and GABA transmission. Simultaneous CA3 and CA1 field potential recordings were obtained during the perfusion of hippocampal slices with the aCSF containing AMPA, NMDA and GABA receptor antagonists. Under these conditions, recurrent field discharges synchronous between CA3 and CA1 were recorded. Field discharges were blocked by addition of calcium-channel blocker Cd^2+^ and disappeared in CA1 after a surgical cut between CA3 and CA1. Cholinergic antagonist mecamylamine abolished CA3-CA1 synchronous field discharges, while antagonists of α7 and α4β2 nAChRs – MLA and DhβE had no effect. Our results suggest that activation of nicotinic acetylcholine receptors is able to sustain CA3-CA1 synchronous epileptiform activity independently from AMPA NMDA and GABA transmission. In addition, mecamylamine but not α7 and α4β2 nAChRs antagonists reduce bicuculline-induced seizure-like activity. The ability of mecamylamine to decrease hippocampal network synchronization might be associated with its therapeutic effects in a wide variety of CNS disorders including addiction, depression and anxiety.

## Introduction

Acetylcholine (ACh) exerts a wide range of neuromodulatory effects in numerous physiological and pathological states [1]. The action of ACh is mediated by two types of receptors: muscarinic (mAChRs) and nicotinic (nAChRs), named after their respective agonists muscarine and nicotine. While the muscarinic type G-protein coupled receptors (GPCRs) mediate a slow metabolic response via second-messenger cascades, the nicotinic type are ligand-gated ion channels that mediate fast cholinergic synaptic transmission [2–3
]. In hippocampus several subtypes of nicotinic (α7, α4β2 α3β4) and muscarinic (M1M4) acetylcholine receptors are widely expressed in pyramidal cells and interneurons at pre- and postsynaptic sites [4–5]. Activation of AChRs has an important role in hippocampal hypersynchronization and pacing of neuronal activity [6–7]. Cholinergic agonist carbachol induces rhythmic oscillations that resemble patterns of epileptiform activity in vitro [8–10].Cholinergic agonist pilocarpine induces status epilepticus in vivo and recently it was shown that administration of pilocarpine causes a 6-fold increase of hippocampal ACh release paralleling the development of tonic seizures [11–13]. High doses of nicotine also induce seizures in animals, and mutations in genes coding for nAChR subunit are associated with seizures in humans [14–16]. Despite these multiple links to epilepsy, the exact function of cholinergic receptors in patterning of hippocampal synchronization remains unclear.

Synchronization of hippocampal fields is primarily mediated by glutamatergic and GABAergic synaptic transmission. Because of that, the influence of endogenous ACh is easily concealed during field potential recordings. The aim of this study was to investigate the ability of cholinergic neuromodulation to sustain hippocampal field synchronization in the absence of GABAergic and glutamatergic transmission. Field potential recordings were obtained in CA3 and CA1 during perfusion of hippocampal slices with aCSF containing AMPA, NMDA and GABA receptor antagonists. Cholinergic antagonists were added to the perfusion solution to study the effect of AChRs activation during hippocampal field synchronization. We also compared the effects of nicotinic antagonist mecamylamine (MEC) and selective α7 and α4β2 nAChRs antagonists on induced hippocampal seizure-like activity.

## Materials and Methods

### Animals

All experimental procedures were performed on Wistar rats according to the guidelines provided by the National Institutes of Health for the humane treatment of animals and approved by the Animal Care Committee of Bogomoletz Institute of Physiology of National Academy of Science of Ukraine. Postnatal day 10-14 rats were deeply anesthetized using sevoflurane and decapitated. Transverse brain slices were prepared according to previously described techniques [17]. Briefly, brains were removed and placed in the ice-cold aCSF of the following composition (in mM): 126 NaCl, 3.5 KCl, 2 CaCl2, 1.3 MgCl_2_, 1.25 NaH_2_PO_4_, 24 NaHCO_3_, 11 D-glucose). The 500μm thick slices were cut using a Vibroslice NVSL (World Precision Instruments, Sarasota, FL). Slices equilibrated at room temperature and constantly oxygenated aCSF for at least two hours before the experiment.

### Induction of epileptiform activity

Synchronous field discharges were induced by perfusion of hippocampal slices with the low-Mg^2+^ aCSF containing AMPA, NMDA and GABA receptor antagonists, which we refer to as “synaptic blockers aCSF”. Synaptic blockers aCSF has the following composition (in mM): 100 NaCl, 5 KCl, 1 CaCl2, 1.25 NaH2PO4, 24 NaHCO3, 11 D-glucose; and 6,7-dinitroquinoxaline-2,3-dione (DNQX 10μM); S,10R)-(+)-5-methyl-10,11-dihydro-5H-dibenzo[a,d]cyclohepten-5,10-imine maleate (MK-801 2 μM); [R-(R*,S*)]-6-(5,6,7,8-tetrahydro-6-methyl-1,3-dioxolo[4,5-g]isoquinolin-5-yl)furo[3,4-e]-1,3-benzodioxol-8(6H)one (bicuculline 10μM).

Nonsynaptic seizure-like activity (SLA) was induced by perfusion of the hippocampal slices with low-Ca^2+^ aCSF of the following composition (in mM): 115 NaCl, 5 KCl, 1 MgCl_2_, 1.25 NaH2_P_O_4_, 24 NaHCO_3_, 11 D-glucose.

Bicuculline (10 μM) and 4-aminopyridine (4-AP, 100μM) were used to induce SLA in the following aCSF (in mM): 125 NaCl, 5 KCl, 1 CaCl2, 1.3 MgCl_2_, 1.25 NaH_2_PO_4_, 11 D-glucose, 24 NaHCO_3_. Chemicals were purchased from Sigma (St. Louis, MO), DNQX, DhβE, d-tubocurarine were obtained from Tocris (Ellisville, MO).

### Extracellular and patch clamp recordings

For extracellular recordings slices were transferred to a submerged recording chamber and perfused with oxygenated aCSF (22-25°C) at a rate of 2-3 ml*min^-1^. Temperature control was performed with the Dual Temperature Controller (TC-144, Warner Instruments). Simultaneous recordings of field potentials were obtained from the CA3 and CA1 pyramidal cell layer with extracellular glass microelectrodes (2–3 MΩ) filled with aCSF. Signals were low-pass filtered (0.5 kHz), amplified using a 2-channel differential amplifier M1800 (A-M Systems, Carlsborg, WA), digitized at 10 kHz using an analog-to-digital converter (NI PCI-6221; National Instruments, Austin, TX).

Patch clamp recordings were performed simultaneously with extracellular recording to investigate the coincidence of postsynaptic currents and field discharges. CA1 pyramidal cells were visually identified with an infrared-differential interference contrast (IR-DIC) microscope (Olympus BX50WI) and captured with a CoolSNAP ES2 (CCD ICX285) video camera. Spontaneous postsynaptic currents were recorded from CA1 pyramidal cells using a patch clamp technique in a whole-cell configuration. Patch electrodes were fabricated from borosilicate glass capillaries of 1.5 mm outer diameter (Sutter Instruments, USA) using a programmable puller (P-97; Sutter Instruments, USA). The recording pipettes were filled with (in mM): 100 Cs-gluconate, 17.5 CsCl, 8 NaCl, 10 HEPES, 10 EGTA, 2 MgATP (pH 7.3). When filled with intracellular solution, recording pipettes typically had resistances of 5–7 MΩ.

### Data analysis

Data were analyzed with WinWCP (Strathclyde Electrophysiology Software, University of Strathclyde, Glasgow, UK), Clampfit (Axon Instruments), Origin 8.0 (OriginLab, Northampton, MA). Cross-correlation analysis was used to determine the level of synchronization between CA3 and CA1 field potential recordings. The sampling data of recordings were filtered by the low pass digital Gaussian filter with a cut-off frequency of 50 Hz. Cross-correlation function (CCF) was then calculated for paired signal samples and smoothed using Lowess smoother (span = 0.01). Next, the first CCF maximum was measured to estimate the level of CA3-CA1 field potential synchronization. Results are reported as mean cross-correlation value ± standart deviation. Summary data are presented as mean ± SD. Two-sample t-test, paired t-test, paired sample Wilcoxon signed-rank test were used for statistical analysis and p < 0.05 was considered statistically significant.

## Results

### CA3-CA1 synchronous field discharges induced in aCSF with AMPA, NMDA and GABA antagonists

Perfusion of hippocampal slices with aCSF containing AMPA, NMDA and GABA antagonists (DNQX 10μM, MK-801 2μM, bicuculline10μM) resulted in the development of robust epileptiform activity in CA3 and CA1 hippocampal areas (Fig 1A). This epileptiform activity represented rhythmic field discharges that fired continuously or were arranged in bursts. (Fig 1B). Field discharges had a mean duration of 1.07 ± 0.34 sec and mean frequency of 0.04 ± 0.02 Hz (n = 20, Fig 1C). Bursts of field discharges had a mean duration of 36.37 ± 11.84 sec and appeared with a mean inter-burst interval of 267.09 ± 146.80 sec; inside of a burst, mean frequency of field discharges was 0.48 ± 0.26 Hz (n = 12, Fig 1B). Field discharges and bursts were synchronized between CA3 and CA1 (cross correlation 0.47 ± 0.17, n = 25). Hippocampal slices are known to produce epileptiform bursting under nonsynaptic conditions such as low-Ca^2+^ milieu [18–19]. However, here we hypothesize that different, synaptic mechanisms account for field discharges synchronization induced in aCSF with AMPA, NMDA, GABA antagonists, unlike nonsynaptic mechanisms of low-Ca^2+^ SLA (Fig 1D). The level of synchronization between CA3 and CA1 in low-Ca^2+^ aCSF (cross-correlation 0.05 ± 0.04, n = 12) was significantly lower (p < 0.001) than synchronization of field discharges induced in aCSF with synaptic blockers (Fig 1).

**Fig. 1.**
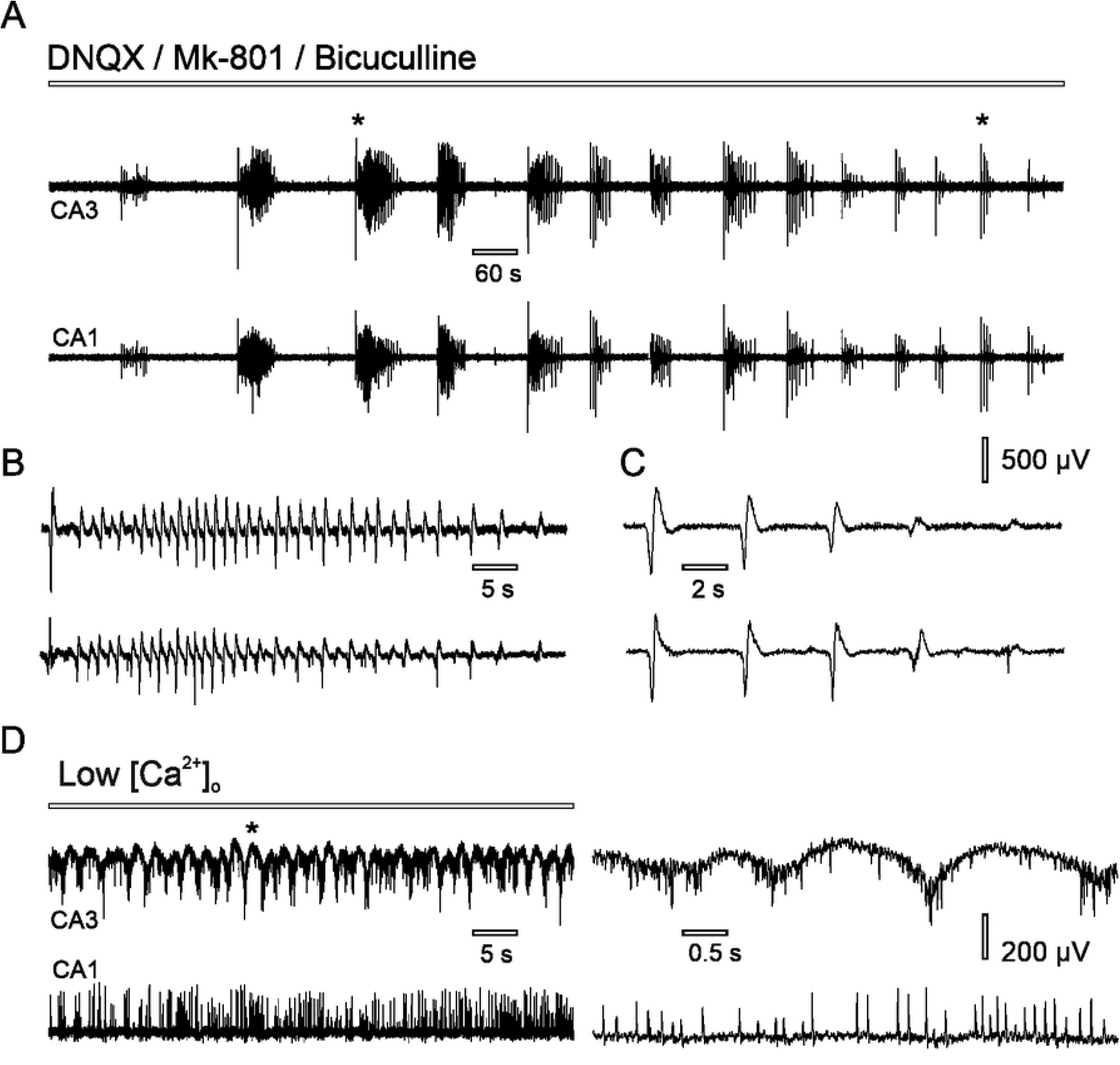
Hippocampal field discharges in CA3 and CA1 under nonsynaptic conditions. (A) Simultaneous recording of synchronous field discharges induced in synaptic blockers aCSF; fragments of the recording marked with asterisk are shown on B and C. (B) Burst of field discharges. (C) Single field discharges. (D) Nonsynaptic population spikes induced in low-Ca^2+^ aCSF; portion of the recording marked with asterisk is shown on the left.

Application of CdCl_2_ abolished synchronous field discharges induced in aCSF with synaptic blockers (Fig 2A). Following 10 min of stable CA3-CA1 synchronous field bursting (cross-correlation 0.44 ± 0.14), 15 μM CdCl_2_ was added to the perfusion aCSF, which resulted in complete blockade of synchronous field discharges (cross-correlation 0.06 ± 0.02, n = 22 p < 0.001). Mean delay time for field discharges abolishment was 2.68 ± 2.45 min (n = 22); prolonged perfusion with CdCl_2_ resulted in the development of nonsynaptic population spikes (n = 10, Fig 2A), which were similar to population spikes induced in low-Ca^2+^ aCSF. Simultaneous patch-clamp and field potential recording during perfusion with aCSF containing AMPA, NMDA and GABA antagonists revealed the coincidence of synaptic currents with synchronous field discharges but not with the nonsynaptic population spikes (Fig 2B). Mechanical separation of CA3 and CA1 hippocampal fields resulted in complete abolishment of field discharges in CA1 but not in CA3 (cross-correlation 0.51 ± 0.21, after the surgical cut – 0.13 ± 0.07, n = 10, p = 0.002, Fig 2C).

**Fig. 2.**
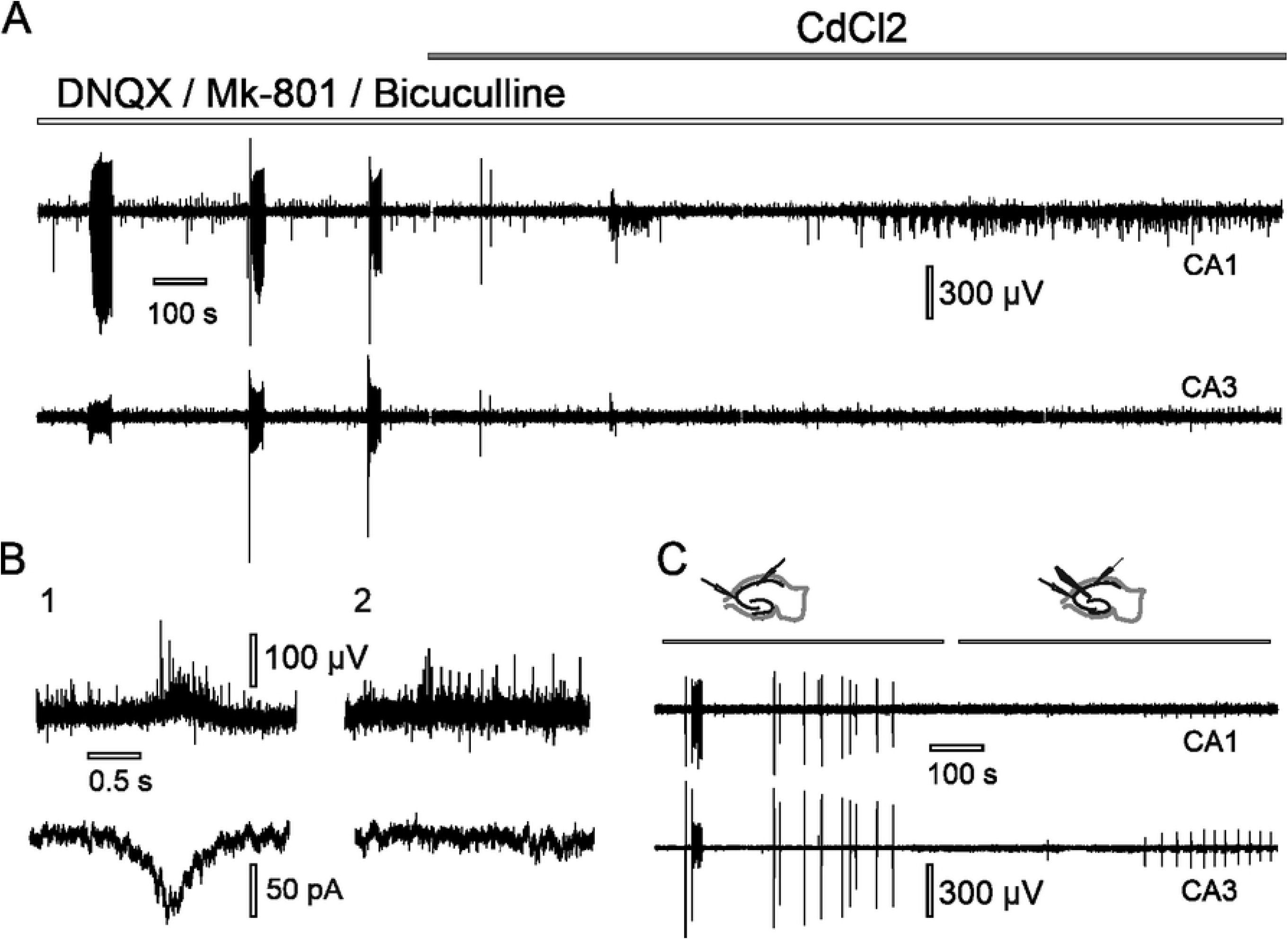
CA3-CA1 synchronization of field discharges induced in aCSF with AMPA, NMDA, GABA antagonists depends on synaptic connections. (A) Blockade of the synchronous field discharges following CdCl_2_ application and the development of nonsynaptic population spikes in CA1. (B) Simultaneous extracellular (upper trace) and intracellular (bottom trace) recording in CA1 during perfusion with synaptic blockers aCSF reveals synaptic currents appear during field discharges (1) but not during nonsynaptic population spikes (2). (C) Field discharges disappear in CA1 but not in CA3 after a surgical cut was made between CA3 and CA1 recording sites.

### Nicotinic acetylcholine receptors account for CA3-CA1 synchronous field discharges induced in aCSF with AMPA, NMDA, GABA antagonists

Synchronous field discharges were blocked following application of nicotinic antagonist d-tubocurarine. Application of muscarinic antagonist atropine did not abolish synchronous field discharges. Application α7 and α4β2 nAChRs antagonists, MLA and DhβE respectively, had no significant effect on CA3-CA1 synchronization (Fig 3C, 3D). Application of nonselective nicotinic antagonist MEC resulted in complete abolishment of field discharges and significant reduction of CA3-CA1 synchronization. Results are presented in Table 1.

**Fig. 3.**
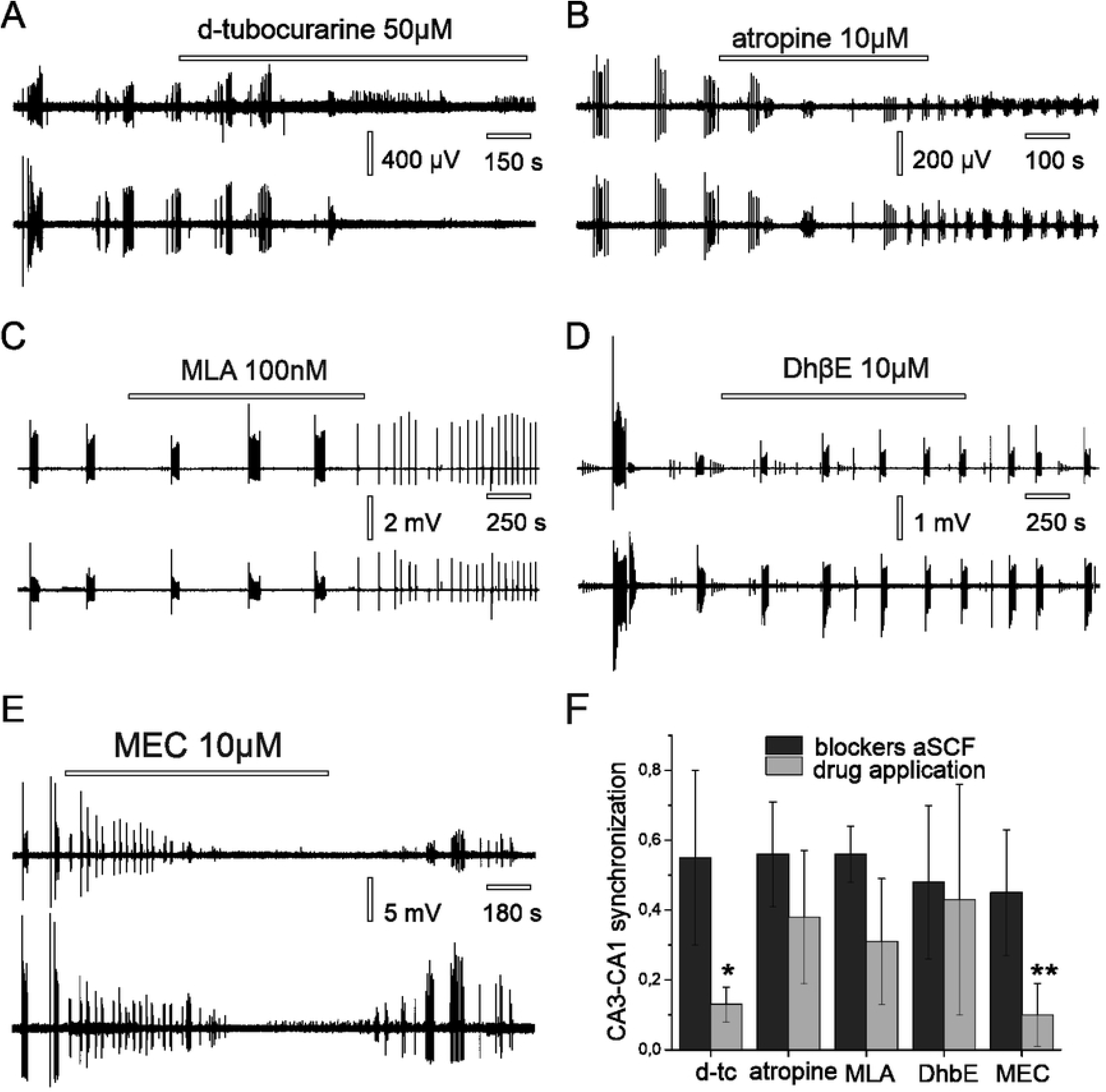
Effect of cholinergic antagonists on CA3-CA1 synchronous field discharges induced in aCSF with AMPA, NMDA and GABA blockers. (A) Application of d-tubocurarine causes reduction of CA3-CA1 synchronous field discharges. (B) Application of muscarinic antagonist atropine does not block CA3-CA1 synchronous field discharges. (C) Antagonist of α7 nAChRs - MLA has no effect on synchronous field discharges. (D) Antagonist of α4β2 nAChRs - DhβE has no effect on synchronous field discharges. (E) Nonselective nicotinic antagonist mecamylamine (MEC) completely abolishes CA3-CA1 synchronous field discharges. (F) Summary data of the effect of cholinergic antagonists on cross-correlation between CA3 and CA1 in synaptic blockers aCSF.

**Table 1.**
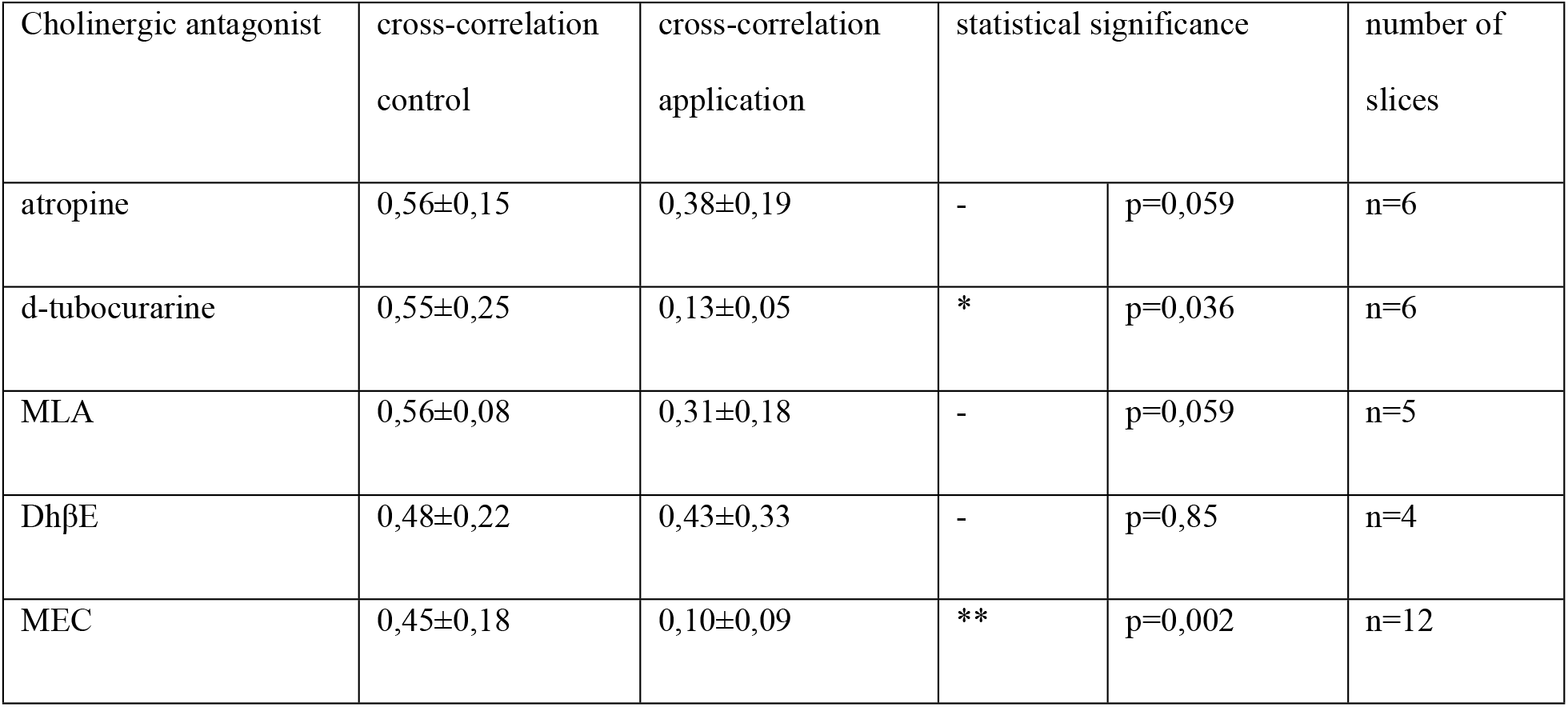
Effect of cholinergic antagonists on CA3-CA1 synchronization of field discharges.

### Effect of MEC on hippocampal SLA induced by bicuculline and 4-AP

Next, we tested MEC on its potential antiseizure properties in two models of SLA: bicuculline and 4-AP. Application of MEC (50 μM) significantly reduced amplitude of bicuculline-induced SLA (in CA3: 2.04 ± 1.04 mV vs 1.61 ± 0.95 mV, n = 10, p = 0.002; in CA1: 3.45 ± 1.87 mV vs 2.19 ± 1.13 mV, n = 10, p = 0.003, Fig 4A). There was no significant effect on the frequency of bicuculline-induced SLA following MEC application (0.13 ± 0.07 Hz vs 0.15 ± 0.08 Hz, n = 10, p=0.27). Application of the selective α7 nAChRs antagonist – MLA (100nM) and selective antagonist for α4β2 nAChRs – DhβE (10μM) had no effect on bicuculline-induced SLA (amplitude in CA3: 6.88 ± 3.04 mV vs 5.7 ± 3.1 mV, n = 3, p=0.25 / in CA1: 5.03 ± 0.66 mV vs 4.38 ± 0.46 mV, n = 3, p = 0.25; frequency: 0.06 ± 0.01 Hz vs 0.06 ± 0.03 Hz, n = 3, p = 1, Fig 4B). Additionally, application of MEC had no significant effect neither on amplitude (in CA3: 3.49 ± 2.09 mV vs 2.93 ± 1.58 mV, n = 5, p = 0.81 / in CA1: 1.47 ± 0.45 mV vs 1.34 ± 0.61 mV, n = 5, p = 0.78), nor on frequency (0.36 ± 0.06 Hz vs 0.33 ± 0.07, n = 5, p = 0.62) of SLA induced by 4-AP (Fig 4C).

**Fig. 4.**
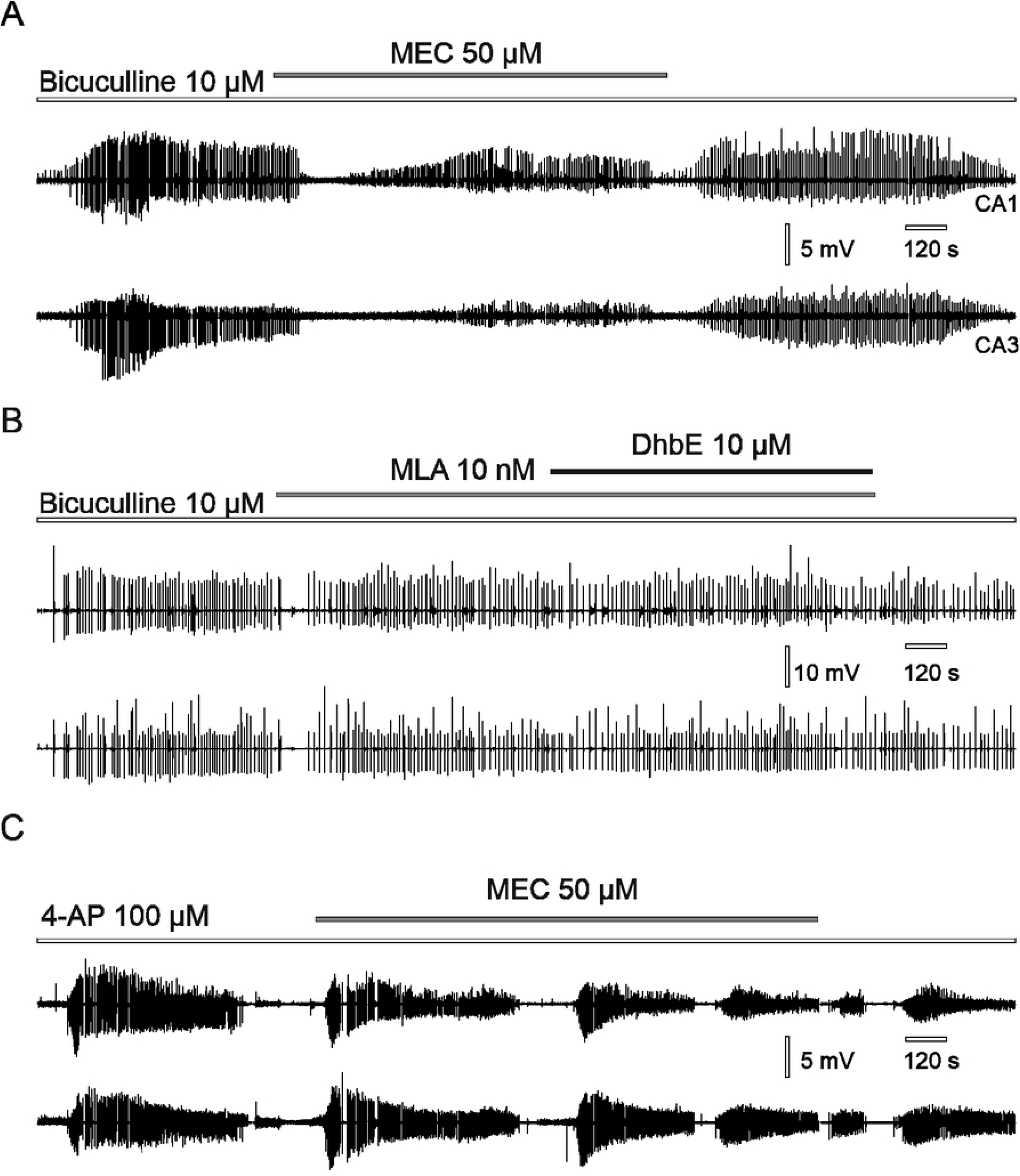
Effect of the cholinergic antagonists on hippocampal SLA activity. (A) Reduction of the bicuculline-evoked SLA following MEC application. (B) Application of α7 nAChRs antagonist MLA and α4β2 antagonist DhβE has no effect on bicuculline-evoked SLA. (C) Application of MEC has no effect on 4-AP induced SLA.

## Discussion

While most of hippocampal synaptic interactions are mediated by glutamate and GABA receptors, neuromodulation through other synaptic systems, such as ACh, exerts powerful effects on network function [21]. Considering the extreme hippocampal propensity for synchronization, we hypothesized that role of endogenous ACh in the hippocampal field potential synchronization might be detected under conditions of increased neuronal excitability and in the absence of AMPA, NMDA and GABA transmission. Since net electrical activity is mostly inhibited under these conditions, we increased neuronal excitability by decreasing osmolarity, omitting Mg^2+^ and increasing K^+^ concentration in perfusion aCSF as described in earlier studies [17–18]. Under these conditions of increased neuronal excitability, we observed synchronous field discharges between CA3 and CA1.

Hippocampal networks can sustain robust SLA under nonsynaptic conditions, such as in the zero-Ca^2+^ milieu [20, 22]. Further, perfusion of hippocampal slices with glutamate and GABA antagonists was shown to induce nonsynaptic bursting similar to low-Ca^2+^ discharges [19]. However, synchronization of discharges between hippocampal areas has never been observed under nonsynaptic conditions. In the present study, we report CA3-CA1 synchronous field discharges in the presence of AMPA, NMDA and GABA antagonists. Addition of CdCl_2_ completely abolished synchronous field discharges, indicating their dependence on voltage-gated calcium channel activation. After mechanical separation of CA1 from CA3, discharges remained unaffected in CA3 but disappeared in CA1, suggesting CA3 as generating site. Simultaneous patch-clamp and field potential recordings of postsynaptic activity revealed coincidence between postsynaptic currents and field discharges. Taken together these results suggest that observed field discharges have synaptic origin.

ACh exerts multiple effects on hippocampal functioning through a wide range of nicotinic and muscarinic receptors [23–25]. In the present study, atropine had no significant effect on synchronization of field discharges. However, perfusion with nicotinic antagonists d-tubocurarine or MEC completely abolished CA3-CA1 synchronous field discharges. These results suggest that activation of nAChRs is able to sustain hippocampal CA3-CA1 synchronization independently of AMPA, NMDA and GABA conductivities.

Three main types of nAChRs are described on hippocampal neurons, namely α7, α4β2, and α3β4 [26–27]. In our experiments, antagonists of α7 and α4β2 nAChRs – MLA and DhβE respectively had no effect on CA3-CA1 filed discharges synchronization. Meanwhile, MEC, a nonselective and noncompetitive nAChRs antagonist, readily abolished CA3-CA1 synchronous field discharges and blocked postsynaptic currents recorded during these events. MEC was initially developed as an antihypertensive medication, but has been studied recently for its therapeutic potential in several neuropathological conditions [28–29]. Beneficial effects of MEC has been reported for epilepsy, substance abuse, depression and anxiety [30–33] and recently it was shown, that MEC reduces levels of ACh and decreases seizures in pilocarpine-induced status epilepticus in rats [11]. Here, we report that MEC has a substantial effect on hippocampal network synchronization, and that this effect is mediated not throughα7 or α4 nAChRs subtypes, implying a possible role for α3β4 subtype. We further studied the effects of MEC on SLA induced by bicuculline and 4-AP. Application of MEC did not change SLA in the 4-AP model, suggesting that nAChRs are not involved in this model of epilepsy. However, MEC caused a significant decrease in the amplitude of bicuculline-induced SLA. Application of α7 and α4β2 antagonists MLA and DhβE had no effect on bicuculline-induced SLA, further suggesting that this effect of MEC on bicuculline-induced bursting is mediated not through α7 or α4β2 nAChRs subtypes. Thus, our results support the ability of MEC to decrease hippocampal network synchronization, which could partially explain therapeutic effects of MEC in a wide variety of CNS disorders.

